# A Refined Analysis of Neanderthal-Introgressed Sequences in Modern Humans with a Complete Reference Genome

**DOI:** 10.1101/2024.08.09.607285

**Authors:** Shen-Ao Liang, Tianxin Ren, Jiayu Zhang, Jiahui He, Xuankai Wang, Xinrui Jiang, Yuan He, Rajiv McCoy, Qiaomei Fu, Joshua M. Akey, Yafei Mao, Lu Chen

## Abstract

**Background:** Leveraging long-read sequencing technologies, the first complete human reference genome, T2T-CHM13, corrects the assembly errors in prior references and addresses the remaining 8% of the genome. While the studies on archaic admixture in modern humans so far have been relying on the GRCh37 reference due to the archaic genome data, the impact of T2T-CHM13 in this field remains unknown.

**Results:** We remapped the sequencing reads of the high-quality Altai Neanderthal and Denisovan genomes onto GRCh38 and T2T-CHM13 respectively. Compared with GRCh37, we found T2T-CHM13 has a significant improvement of read mapping quality in archaic samples. We then applied IBDmix to identify Neanderthal introgressed sequences in 2,504 individuals from 26 geographically diverse populations in different references. We observed different pre-phasing filtering strategies prevalently used in public data can largely impact determination of archaic ancestry, calling for consideration on the choice of filters. We discovered ~51Mb T2T-CHM13 unique Neanderthal sequences, which are predominantly located in regions where the variants distinct between the GRCh38 and T2T-CHM13 assemblies emerge. Besides, we unfolded new instances of population-specific archaic introgression in diverse populations, covering genes involved in metabolism, olfactory-related, and icon-channel. Finally, we integrated the introgressed sequences and adaptive signals with all references into a visualization database website, called ASH (www.arcseqhub.com), to facilitate the utilization of archaic alleles and adaptive signals in human genomics and evolutionary research.

**Conclusions:** Our study refines the detection of archaic variations in modern humans, highlights the importance of T2T-CHM13 reference utility, and provides novel insights into functional consequences of archaic hominin admixture.

## Background

The availability of the Neanderthal and Denisovan sequencing data has enabled the studies in admixture between archaic hominins and modern humans, which subsequently discovered their genetic marks in human genomes [1–6]. All non-Africans possess approximately 2% of the genome inherited from Neanderthals [1, 2, 6–9], while Africans also carry apparent Neanderthal ancestry more than previously reported [10]. Individuals from Oceania have ~2-5% of the genome from Denisovan ancestry, with a small amount of Denisovan sequences also present in Asians [7, 8, 11, 12]. In order to illuminate the functional, phenotypic, and evolutionary consequences of archaic admixture, it is crucial to identify introgressed hominin sequences in modern human genomes. For example, there is substantial evidence that some archaic alleles and haplotypes are adaptive and with high frequency in modern human populations [10, 13–19]. So far, the studies of archaic introgression in modern populations have been relying on the human reference genome GRCh37 [8, 10, 11, 13, 14, 20]. However, the different quality and completeness of other reference genomes (GRCh38 and T2T-CHM13) may bring new horizons to this growing research area [21, 22].

Through the dedicated efforts of the Genome Reference Consortium (GRC), substantial progress has been made from GRCh37 to GRCh38 [22]. Despite these advancements, GRCh38 still contains hundreds of gaps and inaccurately assembled regions [23]. However, in early 2022, the Telomere-to-Telomere consortium achieved a significant breakthrough with the publication of the first complete human genome assembly (T2T-CHM13) [21]. This assembly has effectively addressed repetitive and misassembled regions. The emergence of T2T-CHM13 holds great promise in enhancing our comprehension of human genomics, including segmental duplications, repeats, epigenomics, genetic diversity, large-scale genomic differences, and centromeres [24–28]. Nevertheless, the impact of T2T-CHM13 on archaic introgression patterns in modern human populations remains to be elucidated.

In light of this, the complete and high-quality human reference genome, T2T-CHM13, provides us a unique opportunity to reevaluate and advance our understanding of archaic genetic legacy left into modern populations. We unfolded this mystery by remapping the archaic sequencing reads onto GRCh38 and T2T-CHM13 respectively, and recalling the genetic variants in these archaic genomes. Using a non-modern population reference method, IBDmix, we then identified Neanderthal introgressed sequences in large sets of sequenced data from continental populations. We examined the impact of different filtering criteria on genetic variants utilized in the calling pipeline of archaic introgression. Under the consistent filtering criteria, we made novel discoveries regarding the difference of Neanderthal sequences and adaptive introgression signals in T2T-CHM13 compared to the other references. To make the introgressed sequences and functional inferences easy to access, we launched a database website, called ASH (www.arcseqhub.com), visualizing the archaic segments in the genome and the statistics to facilitate the sharing of resources involving archaic hominin admixture studies.

## Results

### T2T-CHM13 improves read mapping quality in archaic samples

The current data for the archaic high-coverage genomes of the Altai Neanderthal and Denisovan individuals is the GRCh37 version from Prufer et al [2, 6]. To keep the analysis consistent, we downloaded the archaic raw sequencing reads from the previous studies [2, 6], remapped them onto three human reference genomes (GRCh37, GRCh38, and T2T-CHM13), and called the variants with GATK [29] respectively (Fig. 1, see “Methods”). As to the modern human genomes, we used the samples that were sequenced to at least 30× coverage from the 1000 Genomes Project (1KGP) and rephased the VCF data with GRCh38 and T2T-CHM13 references due to the discordant pre-phasing strategies [21, 31] (see “Methods”). We matched the analysis pipeline among the reference genomes as closely as possible so that any major differences would be attributable to the reference genome rather than technical issues in the workflow. After data processing in archaic and modern genomes, we used a non-modern population reference method, IBDmix, to identify the Neanderthal introgressed sequences along the genome and detected population-specific adaptive introgression signals in T2T-CHM13, as seen in the flowchart of our pipeline in this work (Fig. 1).

**Fig. 1.**
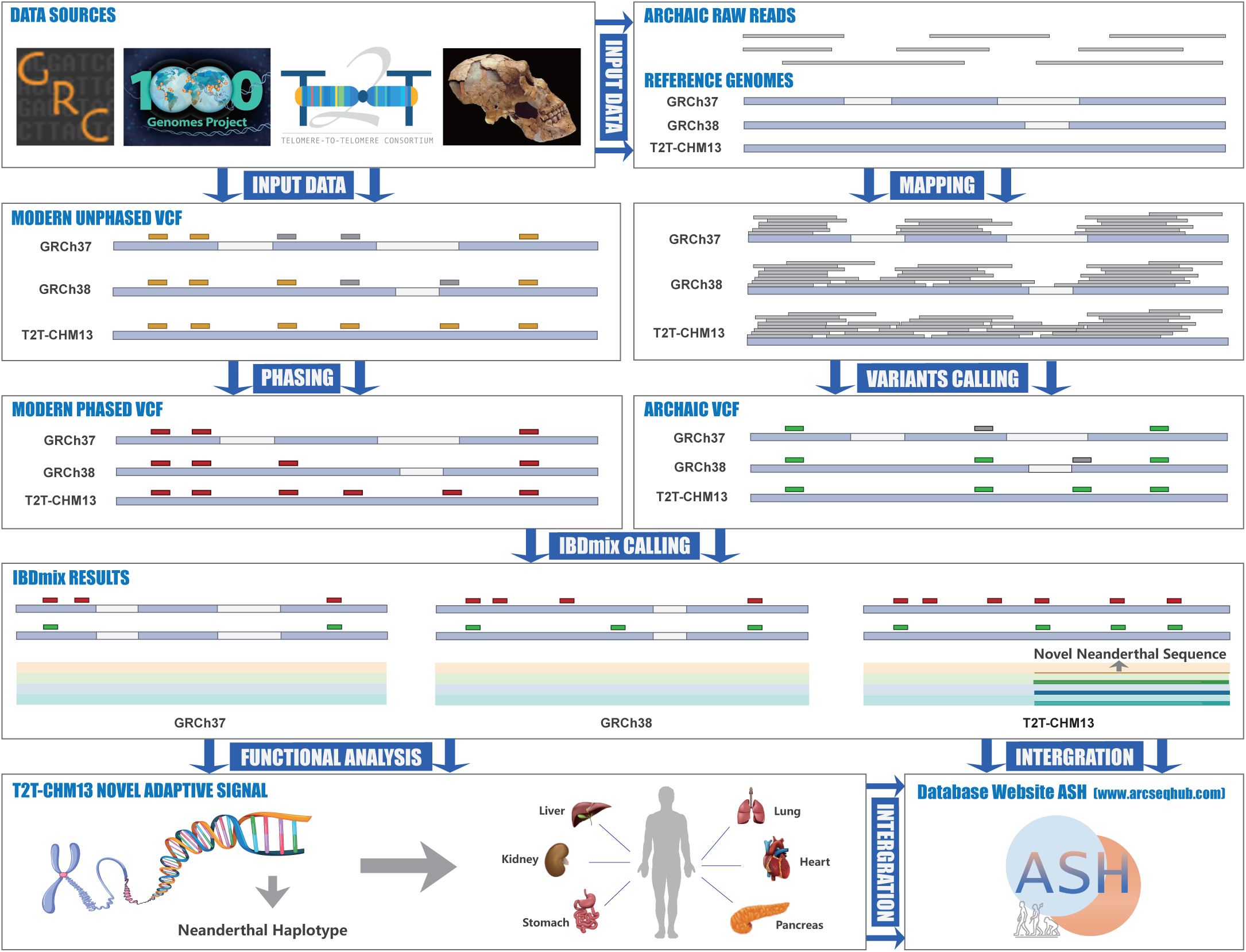
Workflow overview. The core flow of this study includes six steps. (i) Data download. (ii) Rephasing 1KGP unphased VCFs to obtain a phased panel of 2,504 samples. In unphased VCFs rectangles represent high-quality (orange) and low-quality (gray) unphased genotypes. In phased VCFs, red rectangles indicate high-quality phased genotypes. (iii) Remapping archaic sequencing reads on GRCh37, GRCh38, and T2T-CHM13 individually. Rectangles in archaic VCFs represent high-quality archaic variants (green) and low-quality variants (gray). (iv) Detection of archaic introgressed sequence in 1KGP samples based on three references using IBDmix. (v) Functional analysis on adaptive introgression signals in T2T-CHM13. (vi) Integration of the introgression information on three references into a visualization database website, called ASH (www.arcseqhub.com).

Previous study has demonstrated the superiority of T2T-CHM13 as a reference genome for paired-end read alignment in modern human samples across populations [21]. To investigate whether T2T-CHM13 outperforms GRCh37 for short-read alignment in archaic samples, we examined the mapping and calling metrics for the Neanderthal and Denisovan genome data. An additional 1.9 × 10^7^ (1.03%) of sequencing reads were mapped to T2T-CHM13 compared to GRCh38 (Additional file 2: Table S1). Notably, compared to the original GRCh37 reference, T2T-CHM13 has a significant improvement of mapping rate in each chromosome (Fig. 2a). This difference is extremely large for the acrocentric chromosomes, with the rate increasing from ~80% up to > 95%. We also observed a decrease in the standard deviation of the read coverage, a proxy to estimate the mapping quality of complex region [32], in T2T-CHM13 rather than in GRCh38 and GRCh37, indicating the improvement in coverage uniformity with T2T-CHM13 (Fig. 2b). The analysis was replicated in the Denisovan genome data and showed consistent results (Additional file 1: Fig. S1a, Fig. S1b). Overall, these data demonstrate that T2T-CHM13 improves the analysis of read mapping based on the two archaic samples.

**Fig. 2.**
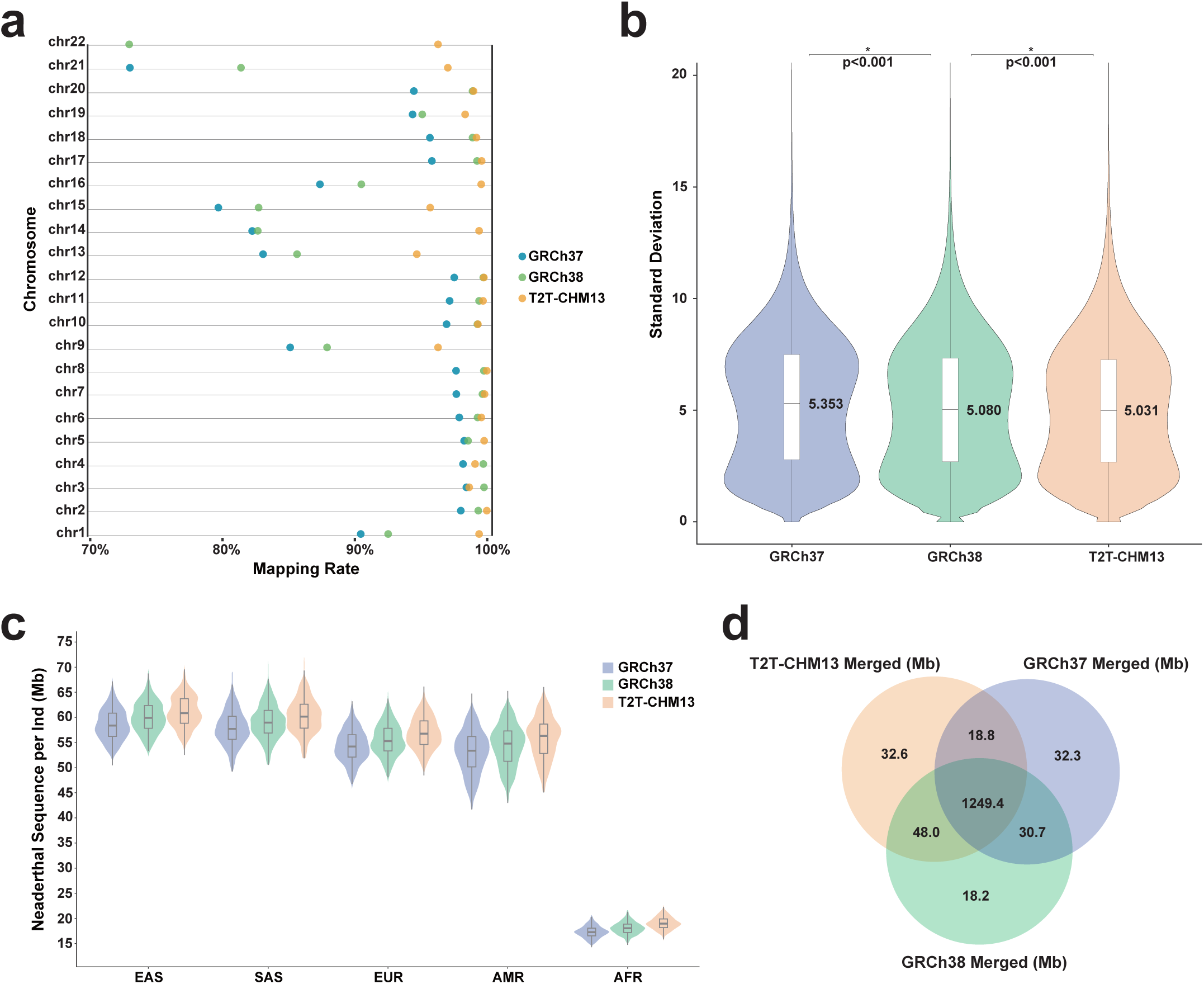
Comparison of Neanderthal ancestry across three reference genomes. **a** Read mapping rate of the Neanderthal sample in GRCh37, GRCh38 and T2T-CHM13. **b** The standard deviation (s.d.) of read count in GRCh37, GRCh38 and T2T-CHM13. **c** Violin plot of Neanderthal sequence identified in five populations from 1KGP data. **d** Venn diagram of the amount of Neanderthal sequence covered in the genome across GRCh37, GRCh38 and T2T-CHM13 references.

### Distinct pre-processing pipelines in public 1KGP phased data bias the proportion of Neanderthal ancestry

As we pre-processed the modern human variation data from 1KGP to prepare for detecting archaic introgression, we noticed considerable discrepancies between the pre-phasing filtering strategies applied to generate the phased Variant Call Format (VCF) files referenced on GRCh38 (Strategy 1) [31] and T2T-CHM13 (Strategy 2) (see “Methods”) [33]. To assess how much these diverse filtering strategies broadly used for public datasets impact the archaic introgression analysis, and also aiming to achieve a universal criteria so that we could minimize the potential differences of archaic sequences attributed to the variations in the reference genome rather than technical pipeline discrepancies, we replicated both strategies on unphased VCFs relative to GRCh38 and T2T-CHM13 respectively [31, 34], utilized Shapeit [35] to do phasing and called Neanderthal sequences using IBDmix [10] with the rephased data.

We observed the number of excluded biallelic variants showed the consistent pattern between GRCh38 and T2T-CHM13 in both pre-phasing strategies, and the main criteria that differed between strategies were the Minor Allele Count (MAC) cutoff and Variant Quality Score Log Odds (VQSLOD) cutoff (Additional file 1: Fig. S2a, Fig. S2b, Additional file 2: Table S2). Notably, singletons are excluded to mitigate potential genotype-calling artifacts in the IBDmix approach utilized for Neanderthal sequence detection. Therefore, much too stringent VQSLOD cutoff implemented in Strategy 2 emerged as a key determinant that potentially introduced bias by producing a larger proportion of evidence against IBD (‘2-0’ genotype pattern variants, see “Methods”), predominantly attributable to the substantial number of variants absent in modern data (Additional file 1: Fig. S3a, Fig. S3b). Factually, we observed the same pattern in the introgressed callsets as expected. In GRCh38 callsets, when using Strategy 1 the mean value of Neanderthal introgressed sequences identified per individual was approximately 15%-20% higher than that when using Strategy 2 (Additional file 1: Fig. S4a). This result is also robust across populations (Additional file 1: Fig. S4a). In addition, when comparing the Neanderthal sequences identified in the callsets relative to T2T-CHM13, we observed the enrichment was even significantly higher in the dataset from Strategy 1, reaching up to ~40% (Additional file 1: Fig. S4b). This result corresponds to our observation with a larger number of ‘2-0’ genotype pattern alleles relative to T2T-CHM13 while using Strategy 2 compared to that in GRCh38 (Additional file 1: Fig. S3a, ~70% versus ~40%).

Collectively, these results uncovered public 1KGP phased panels with different variant pre-processing pipelines may result in significant discrepancies in detecting Neanderthal ancestry by IBDmix. Therefore, in order to strike a balance between the number and the quality of variants required for our analysis of detecting Neanderthal introgression, and also again, to minimize pipeline discrepancies and attribute differences to changes in the reference genome, we performed all subsequent analyses using these high-quality albeit less stringent filtered variants applied by Strategy 1 for datasets relative to GRCh38 and T2T-CHM13.

### IBDmix identified more Neanderthal sequences in T2T-CHM13 reference

After applying the concordant pre-phasing strategy on the datasets of GRCh38 and T2T-CHM13, we identified Neanderthal sequences segregated in modern human populations. All analyses were performed on autosomes and queryable genomic regions only (Additional file 2: Table S3, see “Methods”). The amount of included genomic regions for archaic ancestry detection among reference genomes are rather close (Additional file 2: Table S3). Compared to the callsets in GRCh37 that were previously reported [10], we found more Neanderthal sequences per individual in GRCh38 and T2T-CHM13, which is consistent across populations (Fig. 2C). Although, it is only a modest enrichment with the average amount of ~1 Mb sequences when compared between GRCh37 versus GRCh38 or GRCh38 versus T2T-CHM13 (Additional file 1: Fig. S5, Additional file 2: Table S4). Furthermore, we compared the genomic coverage of Neanderthal sequences among reference genomes, discovered approximately 51.3 Mb of Neanderthal sequences uniquely identified in T2T-CHM13 when compared to GRCh38 (Fig. 2d). Among these novel Neanderthal introgressed regions, approximately 1.68 Mb falls within 8% of the “newly-resolved” sequences in T2T-CHM13 [21]. We also observed a significant overlap of sequences identified among three reference genomes. Of the Neandertal sequences identified in T2T-CHM13, ~94% was shared with the sequences previously reported in GRCh37 (Fig. 2d) [10]. Due to the pipeline discrepancies that GRCh37 is less comparable than the other two references, we focus further analyses on the comparison between GRCh38 and T2T-CHM13 only.

### Small-scale variations impact the identification of Neanderthal sequences

In comparison to GRCh38, we identified a total of 2,087 novel Neanderthal introgressed segments, spanning ~51.3 Mb across the whole genome within the T2T-CHM13 callset. Out of these segments, 242 segments, constituting ~15.92 Mb, do not overlap any introgressed segments identified in GRCh38 and are called “independent sequences”. The remaining 1,845 segments, spanning ~35.35 Mb in total, are extensions of introgressed sequences present in GRCh38 and called “extending sequences” (Fig. 3a). Moreover, the length of these segments exhibits considerable variability, ranging from a few bp to several hundred kb. In the length distribution, we observed the first peak represented the subset of “extending sequences”, while the second peak corresponds to the subset of “independent sequences” (Fig. 3b).

**Fig. 3.**
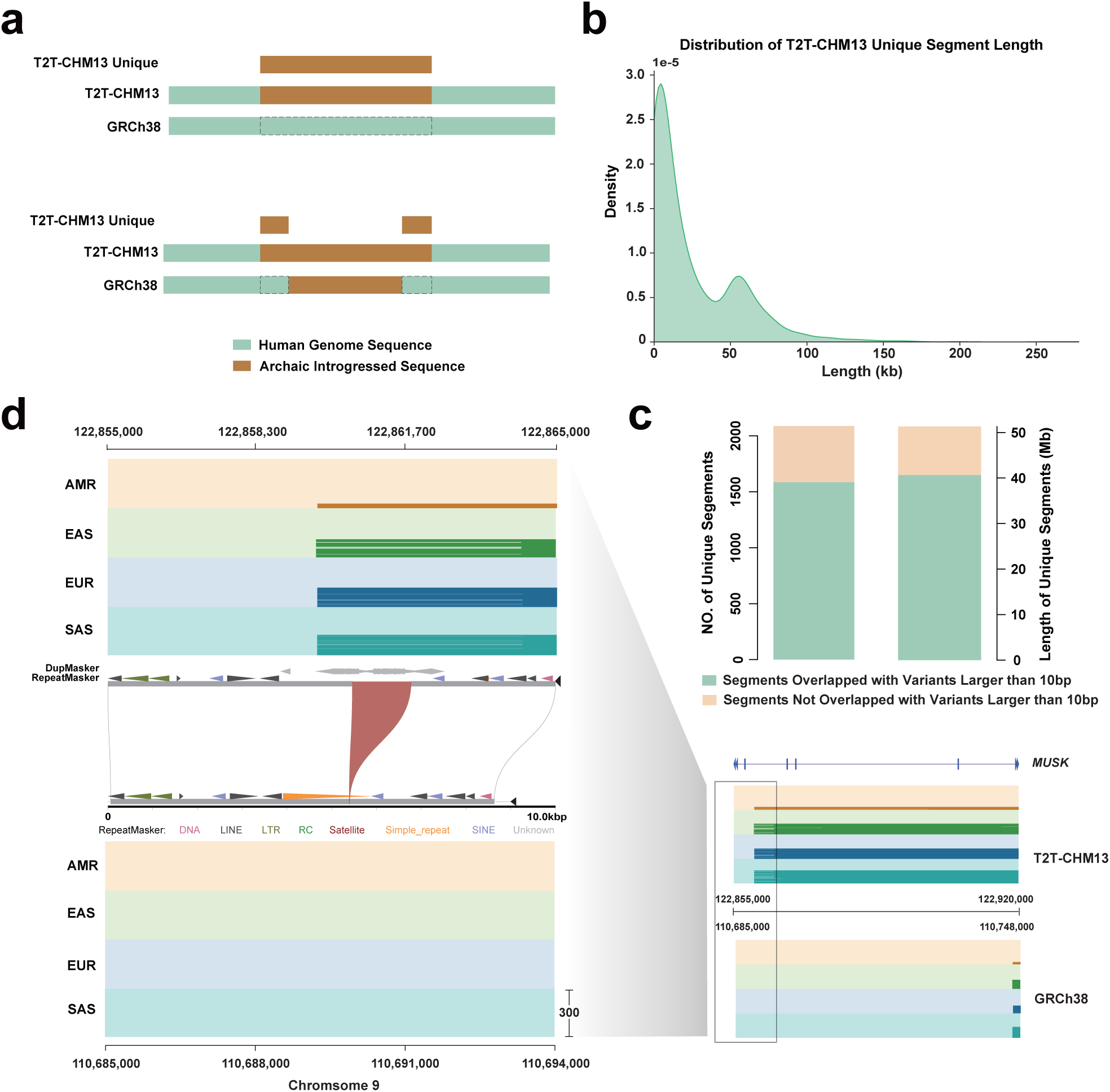
Enrichment of T2T-CHM13 unique introgressed sequences in overlap with genetic variants between T2T-CHM13 and GRCh38. **a** Schematic of T2T-CHM13 unique Neanderthal sequences in the genome. Brown box indicates Neanderthal introgressed sequence, while green box represents modern human sequence. Two types of T2T-CHM13 unique introgressed sequence are shown: completely independent sequences (top) and extensions of the GRCh38 sequences (bottom). **b** Length distribution of T2T-CHM13 unique sequences. **c** Barplot shows the count (left) and length (right) of T2T-CHM13 unique sequences overlap with variants>10bp. **d** An example of T2T-CHM13 unique introgressed sequences covering the *MUSK* gene on chromosome 9. (Inset) The “breakpoint” region of this introgressed sequence. Neanderthal segments are in dark brown (AMR), green (EAS), blue (EUR) and cyan (SAS).

Small differences between T2T-CHM13 and GRCh38 assemblies can impact read mapping quality and genotyping, which may introduce discrepancies in detecting archaic introgressed signals based on IBD inference. In order to examine whether the novel Neanderthal introgressed signals observed in T2T-CHM13 derive from local genetic differences between reference genomes, we leveraged PAV [36, 37] to identify variants across the whole genome between T2T-CHM13 and GRCh38. Subsequently, we systematically screened for local variants with a minimum size of 10 bp, encompassing insertions, deletions, and inversions. We intersected the variants with T2T-CHM13 novel introgressed segments, and found 1,564 segments (74.94% of the total amount of segments), spanning ~40.31 Mb (78.57% of the genomic coverage in total segments) are overlapped with 4,196 variants with length ranging from 10 bp to 1.16 Mb (Fig. 3c, Additional file 2: Table S5).

Furthermore, among the T2T-CHM13 unique Neanderthal introgressed segments that are related to local variants larger than 10 bp, there are intriguing cases that identified novel introgressed segments are commonly shared among populations and enriched with functionally important genes. For instance, a ~1 kb insertion on chromosome 9 in T2T-CHM13 introduces novel signals of Neanderthal introgression, which occur predominantly at high frequency (>5%) in all non-African populations and span the gene MUSK, thought to associate with congenital myasthenic syndrome (Fig. 3d) [38]. This result reveals that local genetic differences between the reference genomes can impact the detection of archaic introgression, whereas the more complete and accurate reference genome T2T-CHM13 can introduce novel knowledge of archaic introgressed signals that were previously undetected.

### Novel signals of adaptive Neanderthal introgression in T2T-CHM13 reference

Previous studies have reported the instances of adaptive archaic introgression, which are the regions where harbor the high-frequency archaic introgressed segments in particular populations [16–19]. To examine the adaptive introgression in alternative reference genomes in addition to discoveries in GRCh37, we followed previously described methods to identify population-level adaptive variants that were located in the introgressed segments called by IBDmix, matched the Neanderthal allele and had large differences in derived allele frequency (DAF) between populations (i.e., Europeans and East Asians, Africans and Europeans, Africans and East Asians) (Fig. 4a, see “Methods”). In total, we identified 87, 87 and 94 population-specific high-frequency Neanderthal haplotypes in GRCh37, GRCh38 and T2T-CHM13, respectively.

**Fig. 4.**
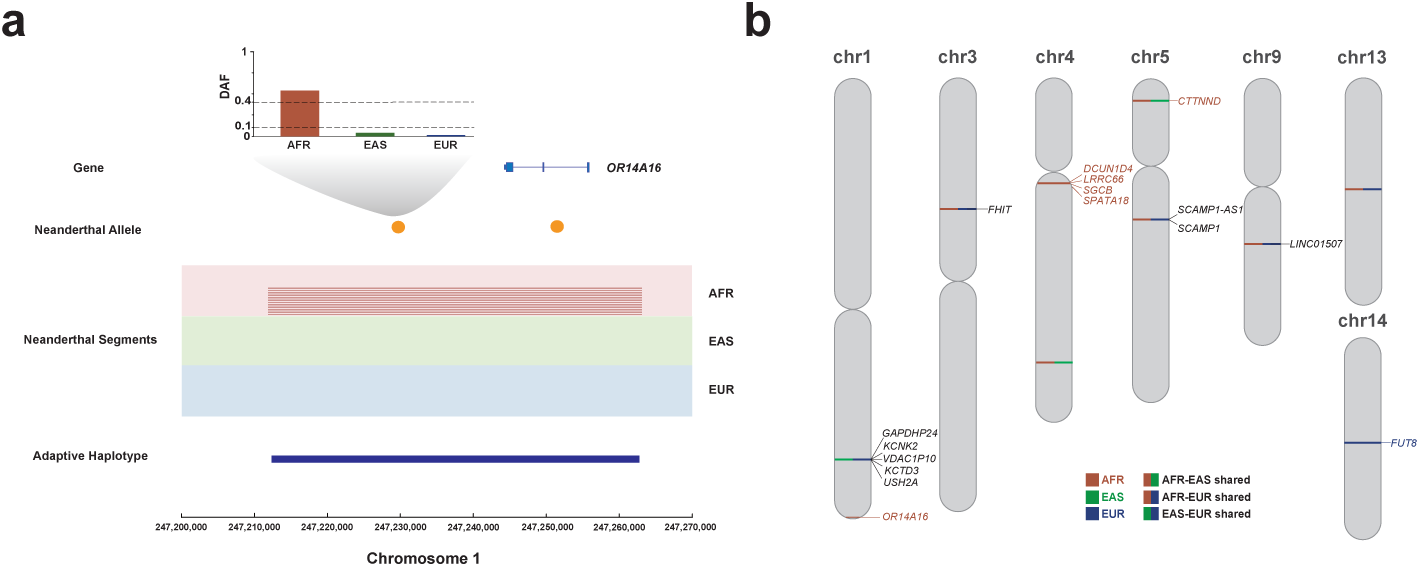
Novel population-specific high frequency introgressed segments in T2T-CHM13. **a** An illustrated example of T2T-CHM13 novel Neanderthal adaptive haplotype in AFR covering the gene *OR14A16* on chromosome 1. AFR-specific high-frequency-derived alleles (DAF>40%) that match the Altai Neanderthal genome are shown as orange circles. Neanderthal segments called by IBDmix are plotted in dark red for AFR, in contrast to the absence of introgression observed in EAS and EUR. The AFR adaptive haplotype merged by Neanderthal segments is shown as dark blue line. **b** Novel population-specific adaptive haplotypes and related genes identified in T2T-CHM13 along the whole genome are shown in AFR (red), EAS (green) and EUR (blue), mixed colors indicate population-shared high-frequency adaptive haplotypes.

We compared high-frequency haplotypes in T2T-CHM13 with those in two other references. Approximately 90% (84/94) of T2T-CHM13 haplotypes were consistent with those found in GRCh37 and GRCh38 results of IBDmix (Additional file 2: Table S6). Of the ten novel population-specific adaptive haplotypes in T2T-CHM13 from IBDmix, 2 were identified as African-specific and 2 were non-African-specific. (Fig. 4b, Additional file 2: Table S7). These regions comprise well-known targets of adaptive introgression such as SGCB, and SPATA18 from previous studies [13, 14, 17]. Notably, these four haplotypes also cover some genomic regions that were for the first time found to be associated with Neanderthal adaptive introgression and showed enrichment for genes involved in metabolism, ion channel, and olfactory function such as FUT8, OR14A16, KCNK2, and KCTD3 (Fig. 4b, Additional file 2: Table S7). In addition, we also identified 4 novel high-frequency haplotypes shared by AFR and EUR and 2 novel high-frequency haplotypes shared by AFR and EAS in T2T-CHM13, which encompass genes enriched in cancer metabolism such as CTNND2, FHIT, and LINC01507 (Fig. 4b, Additional file 2: Table S7). These findings demonstrate that utilizing T2T-CHM13 provides new evidence of adaptive introgression and enhances our interpretation of evolutionary history, variations under selection and with current presence in the human genome.

### Database of the Neanderthal introgressed sequences - ASH

To facilitate more and more widespread application of archaic hominin admixture studies, we developed ArcSeqHub (ASH), a user-friendly interface website. Within this platform, we integrated 985,148, 1,006,918, and 1,037,491 Neanderthal sequences in 2,504 samples from geographically diverse populations (1KGP) and corresponding adaptive Neanderthal introgressed signals in AFR, EAS, and EUR, as identified in GRCh37, GRCh38, and T2T-CHM13, respectively. Utilizing Hypertext Markup Language (HTML) and other scripts, ASH provides users with two distinct search methods: the Gene Query and the Locus Query, which navigates to the visualization of Neanderthal introgressed sequences and related functional genes, along with some statistics such as the introgression ratio across super-populations or individual samples based on three reference genomes (Fig. 5). All statistics and materials presented in ASH are freely downloadable.

**Fig. 5.**
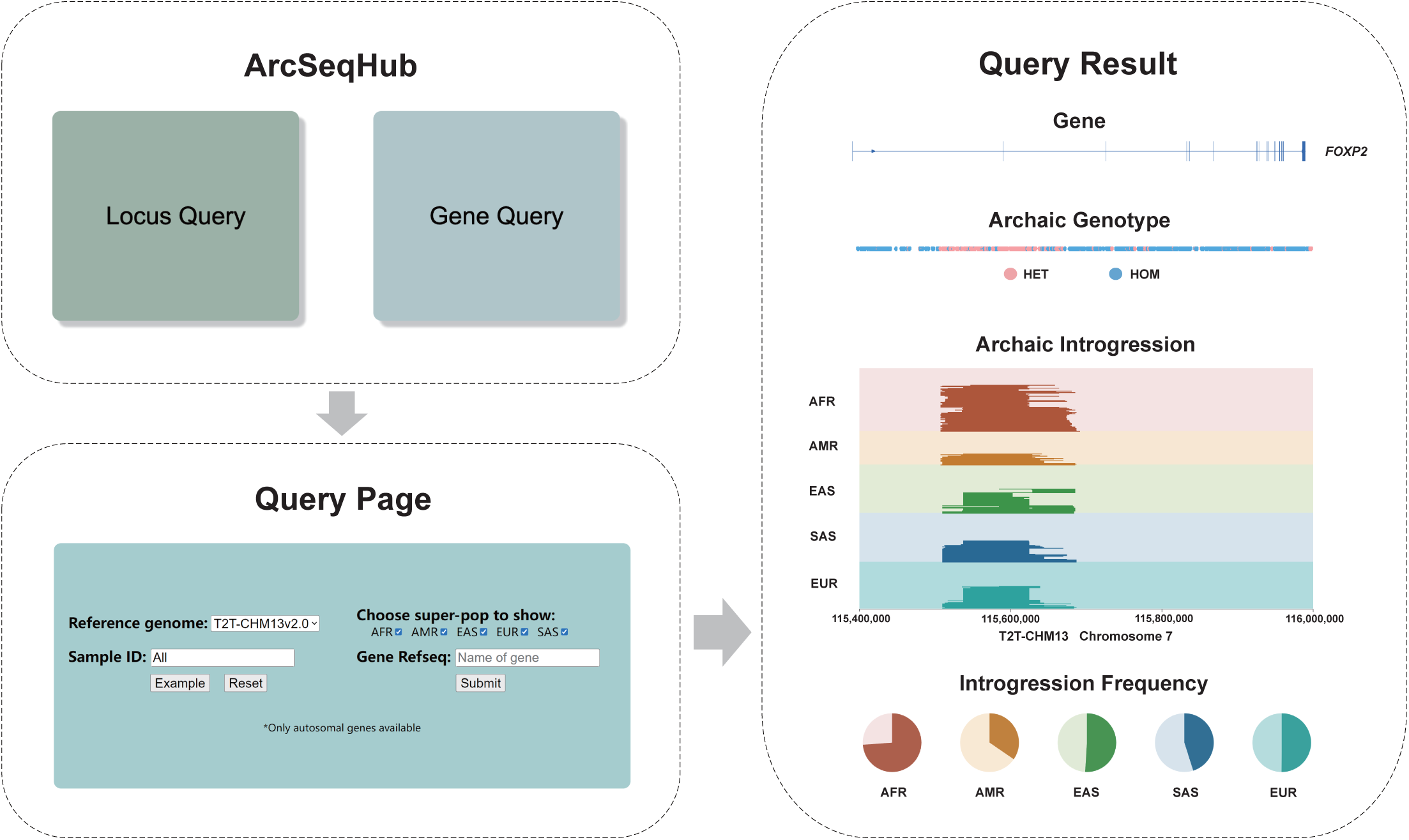
ArcSeqHub (ASH) database website. The information of Neanderthal introgressed sequence and the adaptive signals based on three reference genomes are integrated into a visualization database website (www.arcseqhub.com).

## Discussion

In this study, we realigned and reprocessed the two archaic genomes onto the GRCh38 and T2T-CHM13 references taking advantage of the archaic raw sequencing reads [2, 6]. As such, we were able to make calls and inferences about Neanderthal ancestry in modern populations based on the up-to-date reference genome. We then discovered a modest enrichment of introgressed sequences detected by IBDmix when using T2T-CHM13, compared to other references. We also demonstrate that the improved variant calling in the correctly assembled genome contributes to the unique regions of Neanderthal sequences identified in T2T-CHM13. Furthermore, we updated and refined the population-specific adaptive introgression signals in T2T-CHM13. Besides, we launched a database visualizing archaic introgressed segments in diverse modern populations. Overall, our results provide a valuable resource and novel insights for further archaic admixture studies in T2T-CHM13.

There are several factors predominantly resulting in the difference of archaic sequences detected in T2T-CHM13 versus in other references. One major factor is the improvement of read mapping and variant calling in both archaic and modern human genome data using T2T-CHM13, which would update and correct the genomic position and the genotyping of variants, and thus provide the discrepant information as the key determinant of introgression signal detection. Previous studies have shown that using T2T-CHM13 for read mapping and variant calling enhances the accuracy of modern human genome analysis compared to using GRCh38 and GRCh37 [24]. Similarly, we observed a similar improvement in the analysis of archaic human genomes with T2T-CHM13 (Fig. 2a, Fig. 2b, Additional file 1: Fig. S1a, Fig. S1b). Furthermore, collapsed segmental duplications (SDs) in GRCh38 or GRCh37, such as GPRIN2A/B region [28, 49], can lead to erroneous variant calling, affecting the accuracy of introgressed segment identification. T2T-CHM13 provides a comprehensive SD map, allowing us to exclude these biased regions and resulting in high-quality variant calling for both modern and archaic human genomes. These efforts are crucial for improving downstream introgressed segment detection.

Moreover, we observed that discrepancies in introgressed regions between T2T-CHM13 and GRCh38 often stem from small genetic differences between the two assemblies (Fig. 3c, Fig. 3d). These disparities may result from genomic polymorphisms or inaccuracies present in GRCh38. Such genetic differences can disrupt the signal for initiation and/or extension of introgressed segment detection in IBDmix, thus explaining the observed differences in introgressed sequence identified between GRCh38 and T2T-CHM13. Therefore, besides utilizing a comprehensive genome reference, it is essential to account for reference genetic polymorphisms in future studies of archaic admixture [49]. Furthermore, incorporating a pangenome graph approach into archaic genome studies in the future may offer a promising solution [50].

One more insight afforded by our analyses is that distinct variant filtering pipelines of VCF files can have a significant impact on the amount of Neanderthal ancestry identified in IBDmix. One of the scenarios where there is an strong evidence against the IBD mode (i.e., the allele is less likely to be inherited from the archaic hominins) in IBDmix derives from discordant homozygotes (i.e., variants where the genotype of the archaic individual is homozygous alternative and the modern human genotypes are homozygous reference, referred as ‘2-0’ genotype pattern variants in “Results”) [10]. We especially observed that the more stringent VQSLOD criterion in Strategy 2 excluded a large number of variants. These missing sites would be subsequently regarded as homozygous reference alleles in all modern samples by IBDmix when archaic genotype is homozygous for the alternative allele. This would provide significant evidence against IBD, and lead to the remarkable decrease of Neanderthal sequence. This observation brings up the importance of determining the choice of filters applied on variants prior to the downstream detection of introgressed sequence in archaic admixture studies as the advancements of the genomics era emerge. Researchers often use default program or public-released settings of public data, which is problematic given the default setting might be inappropriate or incompatible with the downstream analyses and thus bias the conclusion [51]. In our analyses, adjusted VQSLOD filters applied in improved reference quality data produce a smaller, more accurate set of variants, while causing more “imputed” sites against IBD according to downstream introgression calling program. This involves trade-offs between variant accuracy and informative variant amount, plus distinct characteristics of methods on detecting archaic ancestry. We emphasize the necessity to pay attention to variant filtering strategies and be cautious to determine the criteria depending on the research question and program algorithm.

In conclusion, we observed the difference of archaic introgressed sequence in T2T-CHM13 detected by IBDmix and demonstrate the rationale behind for such reasons mentioned above. This discrepancy could be also expected when using alternative introgression detection methods based on Hidden Markov Model (HMM) or S* statistics [8, 14, 20]. Nonetheless, it is noteworthy that although we discovered relatively more Neanderthal sequence per individual in T2T-CHM13 by IBDmix, due to the complexity of mixed factors in analyses pipelines and the distinct characteristics of the methods, this consistency of increased amount of archaic sequences in T2T-CHM13 will need to be validated in further studies utilizing alternative methods.

## Conclusions

Taking advantage of the more complete, accurate, and representative reference, T2T-CHM13, we demonstrated the application of the upgraded reference genome plays an important role in the studies of archaic admixture. We found utilizing T2T-CHM13 leads to significant improvement of archaic read mapping. Additionally, the datasets applied with different strategies of pre-phasing filtering that are prevalently used in public data largely showed diverse amount of archaic sequences detected by IBDmix, giving rise to the attention for prudent pre-processing pipeline determination in identifying archaic introgression. Compared to GRCh38, we discovered ~51 Mb T2T-CHM13-unique Neanderthal sequences. In particular, approximately 80% of the unique sequences are overlapped with ~4,200 genetic variants, which are specifically distinct between the T2T-CHM13 and GRCh38 assemblies. Besides, novel population-specific Neanderthal adaptive haplotypes were detected with T2T-CHM13, covering genes involved in metabolism, olfactory-related, and icon-channel, such as *FUT8*, *OR14A16*, and *KCNK2*. Finally, we integrated the Neanderthal sequences in all references into a visualized database website, called ASH (www.arcseqhub.com), to facilitate the usage of archaic alleles and adaptive signals in human genomics and evolutionary research. Together, our work highlights that utilizing the T2T-CHM13 reference can provide novel insights of determining variation in archaic ancestry and further elucidating its functional, phenotypic and evolutionary significance in archaic admixture studies

## Declarations

### Ethics approval and consent to participate

Not applicable.

### Consent for publication

Not applicable.

### Availability of data and materials

The database website ASH can be accessed at www.arcseqhub.com. All data generated or analyzed during this study are included in this published article, its supplementary information files and the database website ASH. The archaic sequencing reads data used in this study can be found in the previous studies [2, 6]. The 1KGP VCF data used in this study can be found in the public database [20, 30, 31].

### Competing interests

The authors declare that they have no competing interests.

### Funding

This work was supported by the National Natural Science Foundation of China (32270668).

### Authors’ contributions

L.C. and Y.M. designed the study. T.R. built up the database website. J.Z., X.W., X.J., Y.H., J.H. and S.L. performed the data analysis. R.M., Q.F., J.M.A., T.R., Y.M., S.L. and L.C. wrote the manuscript. All authors read, edited, and approved the manuscript.

## Acknowledgments

We thank members of the Chen laboratory for their suggestion and discussion. We thank Dr. Aaron B. Wolf for assistance with adaptive introgression analysis.

## Authors’ information

Shen-Ao Liang, Tianxin Ren and Jiayu Zhang contributed equally to this work as first authors.

## Methods and Materials

### Data in this study

We downloaded Altai Neanderthal and Denisovan sequencing reads from http://cdna.eva.mpg.de/neandertal/altai/ [2, 6]. We downloaded 1000 Genomes project phased VCFs phase 3 version 5b from ftp://ftp.1000genomes.ebi.ac.uk/vol1/ftp/release/20130502/ [30]. We downloaded 1000 Genomes 30x genotype VCFs on GRCh38 from http://ftp.1000genomes.ebi.ac.uk/vol1/ftp/data_collections/1000G_2504_high_coverage/working/20201028_3202_raw_GT_with_annot/ [31]. We downloaded 1000 Genomes project genotype VCFs recalled on T2T-CHM13v2.0 from https://s3-us-west-2.amazonaws.com/human-pangenomics/index.html?prefix=T2T/CHM13/assemblies/variants/1000_Genomes_Project/chm13v2.0/ [21]. We downloaded reference genome data from ftp://ftp-trace.ncbi.nih.gov/1000genomes/ftp/technical/reference/human_g1k_v37.fasta.gz (GRCh37), ftp://ftp.ncbi.nlm.nih.gov/genomes/all/GCA/000/001/405/GCA_000001405.15_GRCh38/seqs_for_alignment_pipelines.ucsc_ids/GCA_000001405.15_GRCh38_no_alt_analysis_set.fna.gz (GRCh38) [22] and https://github.com/marbl/CHM13 (T2T-CHM13) [21].

### Mapping archaic sequencing reads

The mapping approaches employed in this study are consistent with those used in previous research [2, 6]. Initially, the first and last two bases were trimmed from the reads to reduce the effects of remaining ancient DNA damage. Then we mapped the reads of Altai Neanderthal and Altai Denisovan to GRCh37, GRCh38 and T2T-CHM13 reference genomes separately using BWA version 0.7.17 [39] with parameters “−n 0.01 −o 2 −l 65536”. Using BWA’s samse command [39], alignments were converted to the SAM format, and then we sorted the SAMs and converted to BAMs with SAMtools version 1.3.1 [40] (coordinate-sorted). Then, we excluded the unmapped single reads, QC-failed single reads, and reads shorter than 35bp. Subsequently, we annotated the NM and MD tag in BAM files using SAMtools calmd [40], and then reads with an edit distance of more than 20% of the sequence length were removed. Finally, we utilized a python script [41] to exclude duplications for each library.

After duplicate removal, we combined the BAM files for all libraries using SAMtools merge [40]. Then we used GATK IndelRealigner [29] to realign sequences in the identified genomic regions. After local realignment, NM/MD fields in BAMs were recalculated using SAMtools calmd [40], and reads with an edit distance of more than 20% of the sequence length were removed.

To obtain mapping statistics for reads, we employed SAMtools stats [40]. To assess the coverage of reads across each base in the genome, we utilized BEDtools version 2.30.0 genomecov [42]. Additionally, to further measure read mapping quality, we calculated the standard deviation (s.d.) of read depth of mapped reads within the specified included region. Initially, contiguous bases in the included regions were treated as individual windows. Then average read depth of each window was estimated, resulting in a set of average read depths for all windows. Subsequently, we obtained the mean read depth across all windows, followed by the calculation of the s.d. of read depth of mapped regions at the whole-genomic level. This approach allows for a comprehensive examination of the distribution of reads across the genome. The utilization of s.d. of read depth of archaic read mapping can be treated as an index to assess the consistency and quality of the mapping process, and the less the read depth s.d. represents the higher the quality of the reference genome.

### Archaic VCF calling

We used the HaplotypeCaller from GATK version 4.3.0.0 [29] to produce genotype calls for single nucleotide variants (SNVs) and insertions and deletions (INDELs) over all sites separately for Altai Neanderthal and Altai Denisovan. To identify high quality variants, we excluded heterozygous variants with aligned reads which carry reference_allele_count/alternative_allele_count less than ⅓ or more than 3. Finally, we generated per-chromosome Variant Call Format (VCF) files in block-gzip compressed form with a tabix (http://samtools.sourceforge.net/tabix.shtml) index file based on three reference genomes respectively.

### Whole genome data processing

Before conducting archaic introgression calling, it is necessary to perform masking operations on the whole-genome data (1000 Genomes, Altai Neanderthal, and Altai Denisovan genomes). Our procedures closely align with those outlined in the previous study [10], and also with some differences in certain details:

#### CpGs mask

CpGs were masked as in [2]. In the case of GRCh38 and T2T-CHM13, we “liftover” the variant data from 15 African hunter-gatherer populations to the GRCh38 and T2T-CHM13 coordinates individually. Different from former studies, when employing the CpG mask from closely related species, for conservative considerations, we included one-to-many alignment scenarios in the generation of sequence alignments for closely related species. Consequently, this allowed for the masking of nearly all CpG sites.

#### Mappability mask

Mappable regions were determined by examining all 35 base long reads that overlap each site. A site is mappable if the majority of overlapping reads are mapped uniquely or without 1-mismatch hit to GRCh37, GRCh38 and T2T-CHM13.

#### Segmental duplication (SD) mask

SD for three reference genomes were removed and downloaded from: https://genome.ucsc.edu/cgi-bin/hgTables [45].

#### Indel mask

Sites within 5 bp of INDELs in VCFs were removed.

#### Accessibility mask

The 1000 Genomes accessibility mask were applied to three reference genomes. GRCh37 accessibility mask data were obtained from: http://ftp.1000genomes.ebi.ac.uk/vol1/ftp/release/20130502/supporting/accessible_genome_masks/20141020.strict_mask.whole_genome.bed

T2T-CHM13 accessibility mask data were obtained from: https://github.com/arangrhie/T2T-HG002Y/tree/main/accessibility_masks

In the case of GRCh38, given the high-coverage nature of the 1000 Genomes data used, the accessibility mask data published was not utilized. Instead, we employed a methodology analogous to that for T2T-CHM13 [24] to generate the accessibility mask for GRCh38.

For archaic samples, we applied three filters as in [2].

#### Tandem repeat filter (TRF)

We downloaded the Tandem Repeat Finder annotation for GRCh37, GRCh38 and T2T-CHM13 from: https://genome.ucsc.edu/cgi-bin/hgTables [45]

#### Mapping quality filter (MQ30)

We computed the root-mean-square mapping quality from BAMs using a custom python script, as opposed to extracting the MQ (Mapping Quality) field from VCFs [2]. This choice was necessitated by modifications in the GATK tools.

#### Genome alignability filter

We produced the map35_50% track, which requires that at least 50% of all possible 35 mers overlapping a position do not find a match to any other position in the genome allowing for up to one mismatch [2].

#### T2T-CHM13 cenSat mask

We merged consecutive centromere/αSat Higher Order Repeat (HOR) array regions until ct (centromeric transition region) appeared in the cenSat annotation track from UCSC, and masked these cenSat regions in T2T-CHM13 archaic introgression analysis.

### Neanderthal introgression calling

Following the completion of all masking procedures, we employed a typical archaic introgression detection method without any modern reference panel, IBDmix [10] to identify Neanderthal sequence in all populations including Africans. Utilizing phased VCFs of 2,504 modern samples, archaic VCFs, and the masked region BEDs, we executed IBDmix with parameters (1) --LOD-threshold 4.0, (2) --minor-allele-count-threshold 1, (3) --archaic-error 0.01, (4) --modern-error-max 0.002, (5) --modern-error-proportion 2 and (6) the cutoff for putative introgressed sequence length at 50 kb. Finally, we obtained five distinct sets of IBDmix results corresponding to three different reference genomes (one for GRCh37, and two datasets each for GRCh38 and T2T-CHM13, owing to distinct pre-phasing filtering strategies).

### Phasing modern VCFs

We needed modern phased 1KGP panels combined with archaic genotype panels to call archaic introgression. However, the pre-phasing filtering strategies applied to the 1KGP phased VCF datasets for GRCh38 and T2T-CHM13 differ [31, 33]. The pre-phasing filtering strategy for public GRCh38 phased data (Strategy 1): (1) FILTER (column in the VCF) = PASS, (2) GT missingness rate <5%, (3) HWE exact test p value > 1e-10 in at least > one super-population, (3) mendelian error rate (MER) ≤ 5%, (4) minor allele count (MAC) ≥ 2. The pre-phasing filtering strategy for public T2T-CHM13 phased data (Strategy 2): (1) exclude FILTER (column in the VCF) = PASS, (2) exclude variants with an alt allele of ‘*’ after multiallelic splitting, (3) exclude GT missingness rate < 5%, (4) exclude Hardy-Weinberg p-value < 1e−10 in any 1000G subpopulation, (5) exclude sites where Mendelian Error Rate (Mendelian errors/num alleles) >= 0.05, (6) exclude homoallelic sites (MAC=0), (7) exclude variants with a high chance of being errors as predicted by computational modeling.

Therefore, to ensure comparability of results, we reapplied the two distinct pre-phasing filtering strategies to the unphased VCF datasets of GRCh38 and T2T-CHM13 separately. Initially, we annotated the unphased VCF files with p values from the Hardy-Weinberg equilibrium (HWE) exact test [43], stratified by super-population, employing the BCFtools version 1.9 fill-tags plugin [44]. Subsequently, multiallelic sites were segregated into distinct rows, and INDELs underwent left-normalized representation using the BCFtools norm tool [44]. Following this, we computed the Mendelian Error Rate (MER) for each variant row using the Mendellian plugin in BCFtools [44]. Finally, we employed the BCFtools toolkit [44] to filter the variants based on predefined two pre-phasing filtering criteria.

After completing the filtering steps, we employed Shapeit version 5.1.1 [35] for variant phasing. We utilized the phase_common module for variants with a minor allele frequency (MAF) greater than 0.1%, generating a haplotype scaffold. Subsequently, phase_rare module was applied for phasing variants with MAF less than 0.1%. Following this, multiallelic sites were merged into separate rows, and the left-normalized representation of INDELs was performed using the BCFtools norm tool [44]. Subsequent to these preprocessing steps, we utilized the BCFtools toolkit [44] to extract 2,504 unrelated individuals, obtaining four phased VCF datasets (two pre-phasing filtering strategies for two reference genomes) containing haplotypes for these 2,504 individuals.

### ‘2-0’ genotype pattern variants

There is an strong evidence against the IBD mode (i.e., the allele is less likely to be inherited from the archaic hominins) in IBDmix derives from discordant homozygotes (i.e., variants where the genotype of the archaic individual is homozygous alternative and the modern human genotypes are all homozygous reference, called as ‘2-0’ genotype pattern variants), which may come from ‘ impute’ as homozygous reference for missing sites in modern humans as there would be no variation at these sites in the modern human panel. Therefore, there are a large number of variants excluded due to much too stringent VQSLOD cutoff in Strategy 2, which may lead to an unexpected larger amount of ‘2-0’ genotype pattern variants and significant bias to detect archaic introgressed signals when combining modern human phased panels with archaic genotype panels.

### T2T-CHM13-unique introgression analysis

Due to the limitations in coverage of modern human data for GRCh37 reference genome and inherent disparities during data processing between GRCh37 and the other two reference genomes, to investigate the implications of reference genome updates on the emergence of novel archaic introgression signals, our analysis focused exclusively on T2T-CHM13 and GRCh38 datasets to mitigate potential biases stemming from data and technical variations.

To identify the novel archaic introgressed sequences in T2T-CHM13, we “liftover” GRCh38 merged callset to T2T-CHM13 coordinates, and the sequences were then excluded from T2T-CHM13’s merged callset. To further scrutinize whether the novel introgression were in those localized different regions between the two reference genomes, we used T2T-CHM13 as the reference genome and GRCh38 as the query to run PAV version 2.0.0 [36, 37], and obtained all variants in GRCh38 relative to T2T-CHM13. Subsequently, we extracted all the variants larger than 10 bp that we think may influence read mapping quality.

To identify the variants overlapped with the T2T-CHM13 novel archaic introgressed sequences, We processed sequences less than 50 kb (Fig. 3a) in a special manner. Considering the IBDmix cutoff for a minimum sequence length of 50 kb, and any alteration in signals within dynamic programming algorithms might hinder the extension of local introgression sequences, therefore, we extended novel archaic introgressed sequences in T2T-CHM13 less than 50 kb and closely connected downstream to GRCh38 introgression sequences, downstream to 50 kb. Similarly, we extended sequences less than 50kb and closely connected upstream to GRCh38 introgressed sequences, upstream to 50 kb. Additionally, for sequences less than 50 kb and tightly connected both upstream and downstream to GRCh38 sequences, we extended them in both directions until reaching 50 kb.

Subsequently, we identified all T2T-CHM13 introgression sequences covering variants larger than 10bp utilizing BEDtools [42]. To identify significant and larger updates in archaic introgressed signals specific to the T2T-CHM13 reference genome, we further required that the novel introgressed sequences must cover structural variations (variants >50 bp), and these sequences had to be present in at least 5% of the total population across a minimum of two modern human populations.

### Identifying novel population-specific adaptive introgressed signals in T2T-CHM13

Initially, we identified population-specific high-frequency introgressed alleles and merged haplotypes for EUR, EAS and AFR based on GRCh37, GRCh38 and T2T-CHM13 reference genomes separately, primarily drawing from the methodology outlined in [10]. As for the infer for derived and ancestral state, in the case of GRCh37, we used derived allele frequencies calculated and ancestral states tagged by 1000 Genomes Project. For GRCh38, the derived and ancestral states were inferred based on the chimpanzee state in the Ensembl v110 EPO 10 primate alignment [46]. As for T2T-CHM13, we “liftover” the GRCh38 data to T2T-CHM13 coordinates, as there is currently no published EPO version available for T2T-CHM13 coordinates.

To identify novel population-specific adaptive signals in T2T-CHM13, we “liftover” population-specific high-frequency introgressed alleles and merged haplotypes previously identified in GRCh37 and GRCh38 to T2T-CHM13 coordinates. Subsequently, we excluded GRCh37 and GRCh38 data with T2T-CHM13 coordinates from T2T-CHM13 callset to obtain the novel introgressed haplotypes set and the novel introgressed Neanderthal alleles set separately employing BEDtools [42]. Finally, we intersected the two datasets utilizing BEDtools [42] to identify novel population-specific merged introgressed haplotypes covered by at least one Neanderthal derived allele.

### The Archaic Sequence Hub website development

The Archaic Sequence Hub (ArcSeqHub) provides a user-friendly interface for efficiently exploring archaic introgression in modern humans. The query page of ArcSeqHub features two approaches: one based on gene name and the other utilizing segment coordinates. Each query allows users to select a subset of super-populations of interest or specify particular samples. The website’s interface was crafted and supported with Hypertext Markup Language (HTML), Cascading Style Sheets (CSS), JavaScript (JS), and Django version 4.1 [47]. In addition, a custom R script was developed in-house to show archaic introgression for human samples with the R package transPlotR [48]. The pie charts were generated with ECharts and enhanced with Bootstrap version 3.4.1. The website operation is supported by the Aliyun server, in conjunction with nginx version 1.20.1. Functional testing has been conducted on various popular web browsers, here, we recommend using Google Chrome for a better experience. We welcome user contributions, suggestions for improvements and bug reports through www.arcseqhub.com/contact/.

## Additional file information

**Fig. S1.**
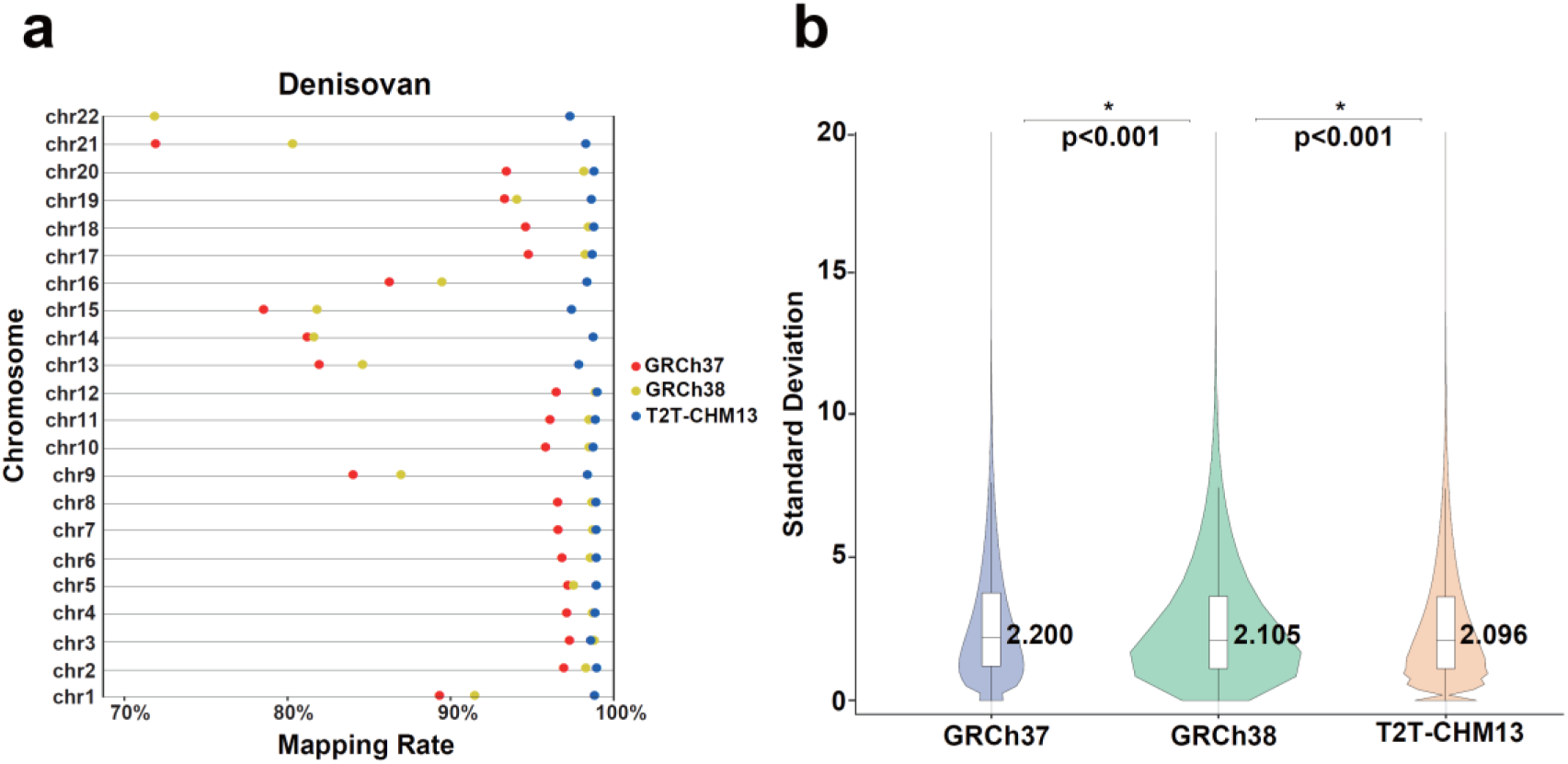
Comparison of Denisovan statistics across three reference genomes. (a) Denisovan read mapping rate in unmasked genomic regions among GRCh37, GRCh38 and T2T-CHM13. (b) The standard deviation (s.d.) of Denisovan read counts in unmasked genomic regions among GRCh37, GRCh38 and T2T-CHM13.

**Fig. S2.**
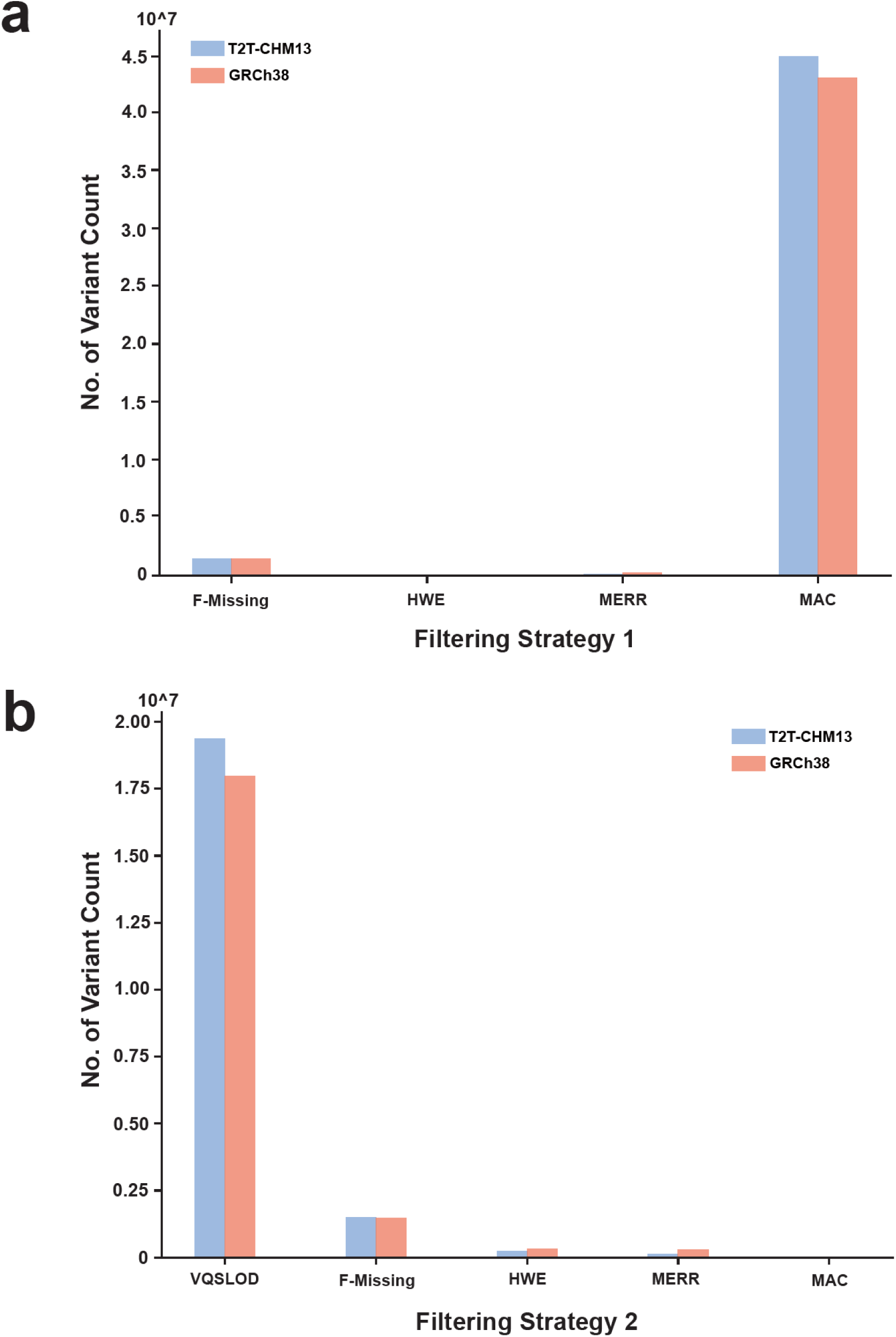
Count of biallelic variants excluded by each filtering criterion in pre-phasing filtering strategies. Bar plots show count of biallelic variants excluded by each filtering criterion in two pre-phasing filtering strategies in T2T-CHM13 and GRCh38. (a) Strategy 1. (b) Strategy 2.

**Fig. S3.**
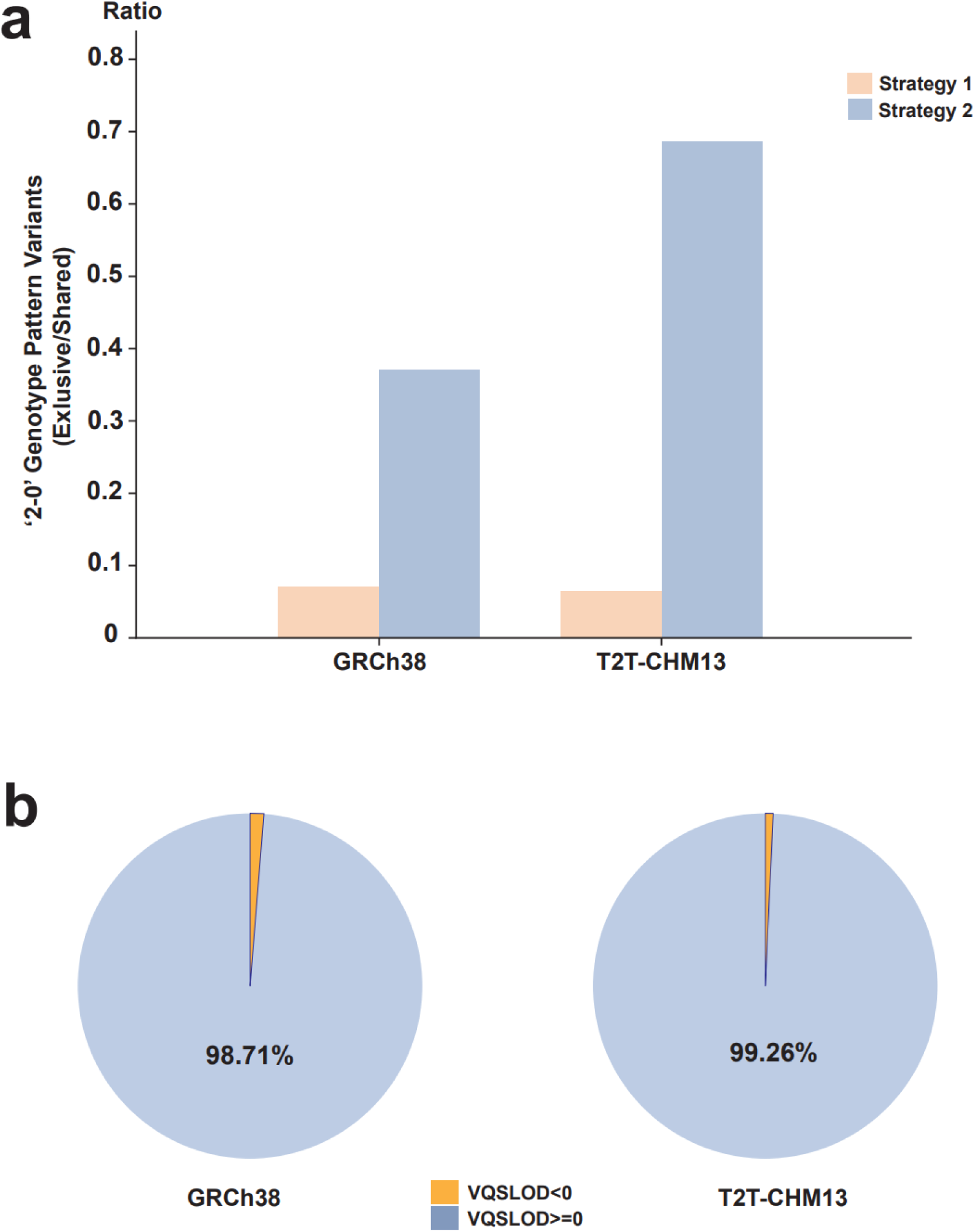
‘2-0’ genotype pattern variants under two pre-phasing strategies. (a) The ratio of exclusive ‘2-0’ genotype pattern variant count under each strategy and two-strategies-shared ‘2-0’ genotype pattern variant count in GRCh38 and T2T-CHM13. (b) Pie charts show the ratio of the ‘2-0’ genotype pattern variants caused by VQSLOD criterion of Strategy 2 in GRCh38 (left) and T2T-CHM13 (right).

**Fig. S4.**
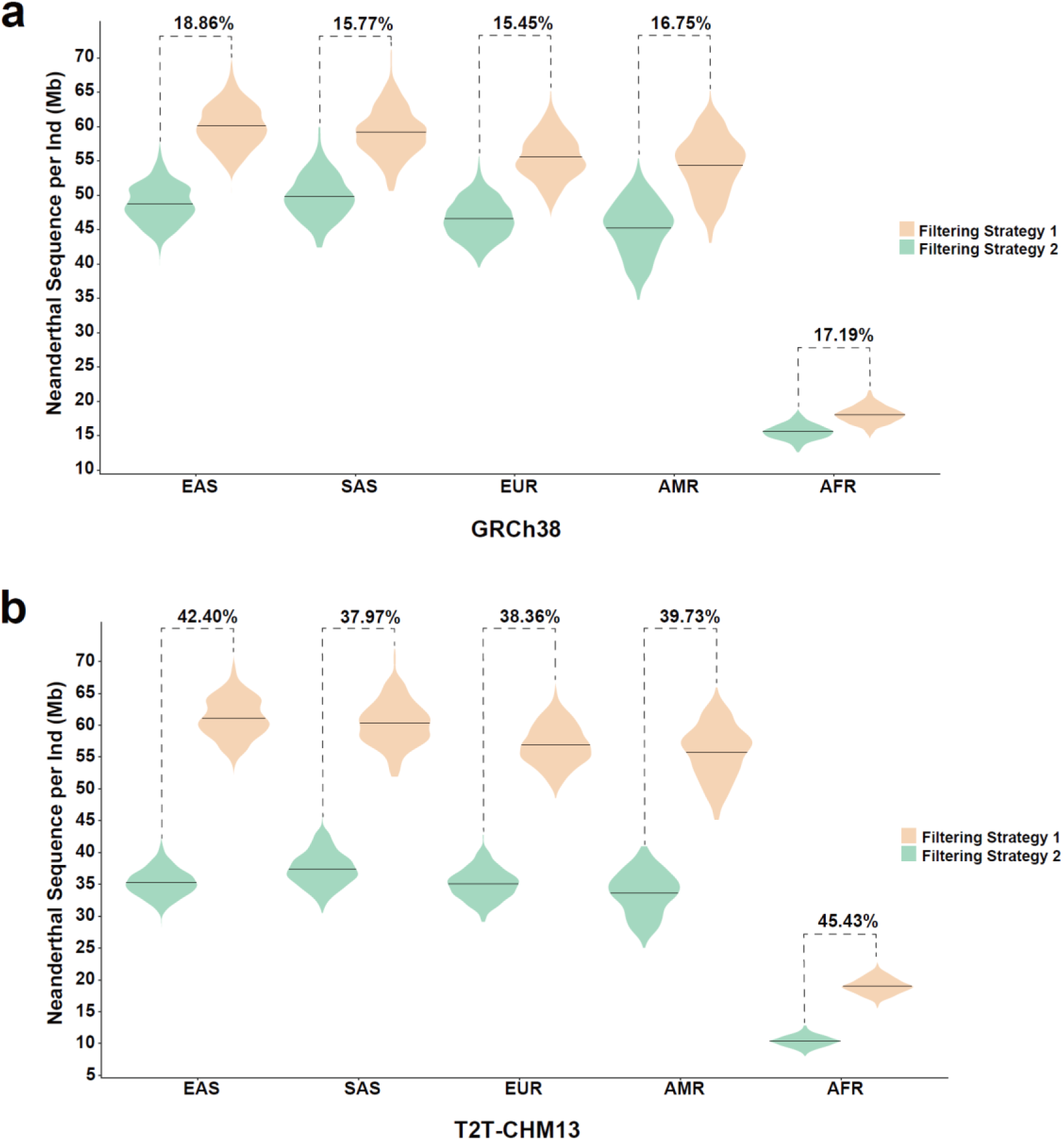
Comparison of Neanderthal introgression detected under different pre-phasing filtering strategies. Violin plots show Neanderthal introgression bias per individual under two different pre-phasing filtering strategies in (a) GRCh38 and (b) T2T-CHM13.

**Fig. S5.**
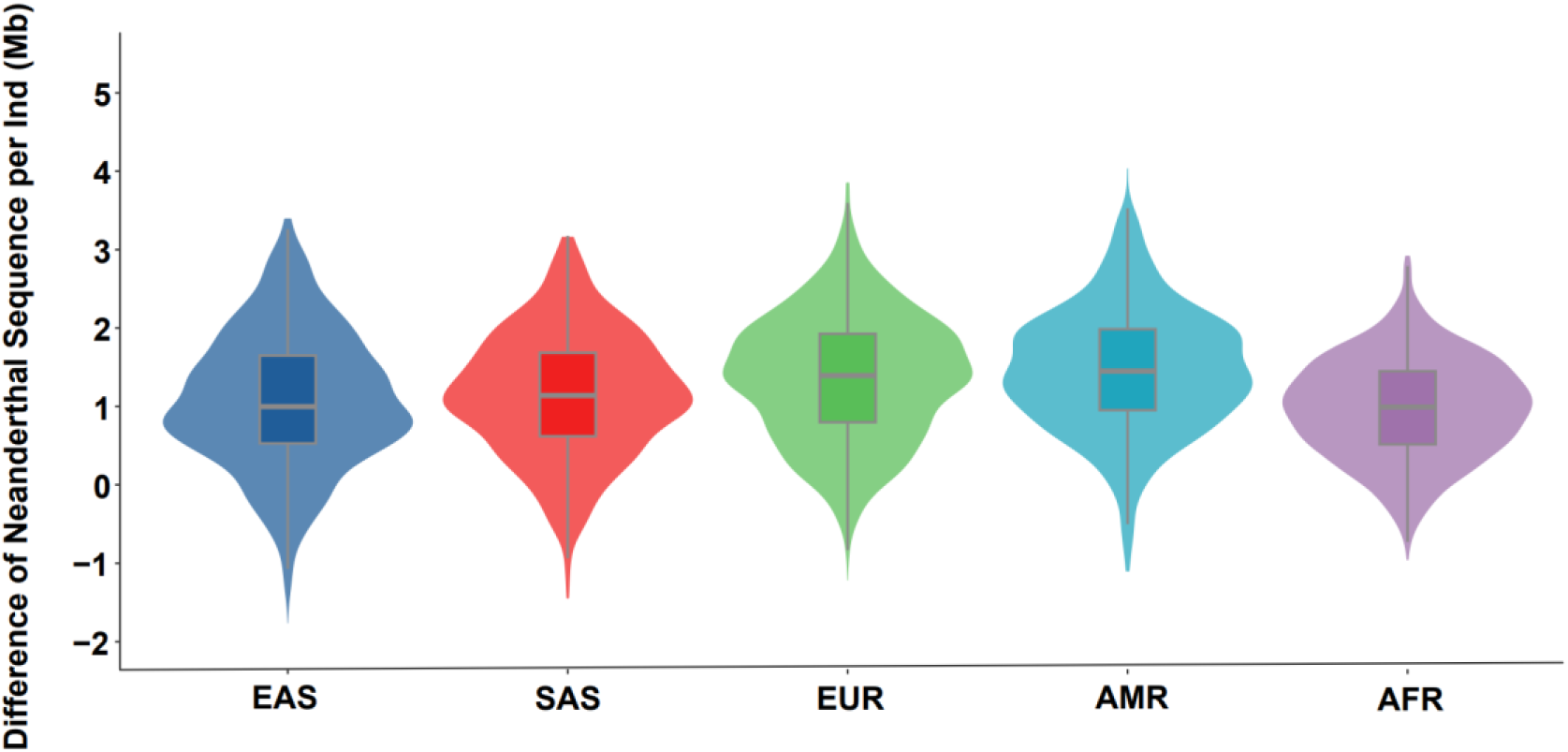
Difference of Neanderthal introgression per individual between T2T-CHM13 and GRCh38. Violin plots show difference (Mb) of Neanderthal introgressed sequences per individual identified between T2T-CHM13 and GRCh38 in five populations from the 1000 Genomes Project (1KGP).

## Additional file 2 File format: XLS

**Table S1. Read mapping statistics of Altai Neanderthal and Denisovan in three reference genomes.** Table shows the count of archaic mapped reads, the total base count of archaic mapped reads (Gb), the error rate of archaic mapped reads and the length (Mb) and mapped rate of genome covered by archaic reads in whole genome level, autosome level and included region.

**Table S2. Count of variants excluded by pre-phasing filtering.** Table shows the count of variants excluded by each filtering criterion in pre-phasing filtering Strategy 1 and Strategy 2 in T2T-CHM13 and GRCh38.

**Table S3. Masked region statistics for Altai Neanderthal and Denisovan in three reference genomes.** Table shows the size (Mb) of included mask, excluded mask, and final included region in Altai Neanderthal and Denisovan in GRCh37, GRCh38 and T2T-CHM13.

**Table S4. Mean value of Neanderthal introgressed sequence in populations in three reference genomes.** Table shows the mean value (Mb/ind) of Neanderthal introgressed sequence identified in 5 populations and 26 subgroups from the 1000 Genomes Project in GRCh37, GRCh38 and T2T-CHM13.

**Table S5. 1,564 T2T-CHM13-unique Neanderthal sequences are overlapped by 4,196 variants larger than 10 bp.** Table shows all of 1,564 T2T-CHM13-unique intervals and intervals expanded to 50 kb are covered by 4,196 variants larger than 10 bp.

**Table S6. Population-specific haplotypes identified in three reference genomes.** Table shows population-specific high-frequency adaptive haplotypes identified based on GRCh37, GRCh38 and T2T-CHM13 reference genomes respectively.

**Table S7. T2T-CHM13-unique population-specific haplotypes and covered genes compared with GRCh37.** Table shows T2T-CHM13-novel population-specific high-frequency adaptive haplotypes and related genes identified in AFR, EAS and EUR compared with GRCh37.

## Notes

http://www.arcseqhub.com

### Competing Interest Statement

The authors have declared no competing interest.

