## Additional file 1 for "A Refined Analysis of Neanderthal-Introgressed Sequences in Modern Humans with a Complete Reference Genome"

### A Refined Analysis of the Neanderthal Sequence in Modern Humans with the Complete Reference Genome

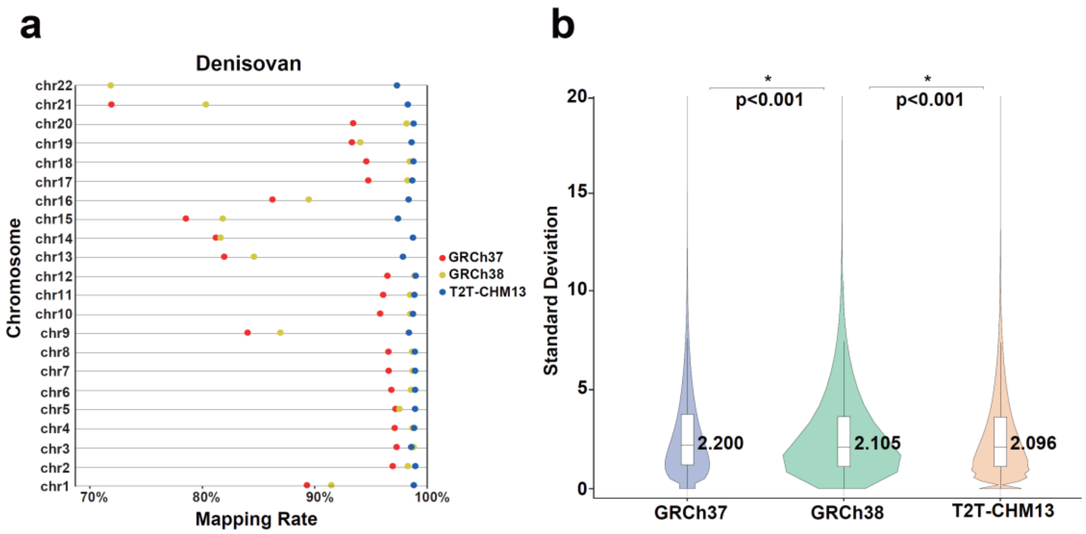


**Fig. S1. Comparison of Denisovan statistics across three reference genomes.** (a) Denisovan read mapping rate in unmasked genomic regions among GRCh37, GRCh38 and T2T-CHM13. (b) The standard deviation (s.d.) of Denisovan read counts in unmasked genomic regions among GRCh37, GRCh38 and T2T-CHM13.

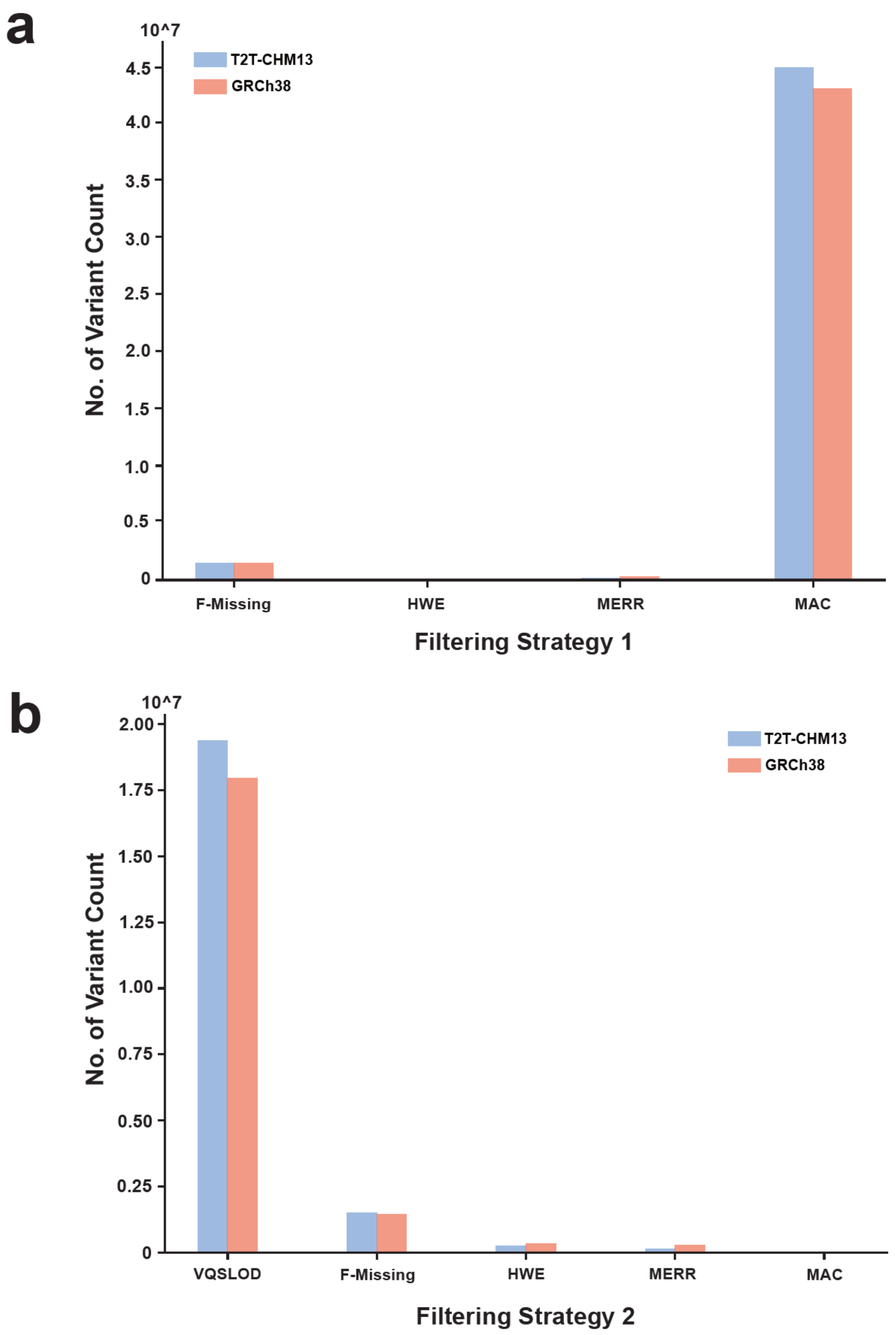


**Fig. S2. Count of biallelic variants excluded by each filtering criterion in pre-phasing filtering strategies.** Bar plots show count of biallelic variants excluded by each filtering criterion in two pre-phasing filtering strategies in T2T-CHM13 and GRCh38. (a) Strategy 1. (b) Strategy 2.

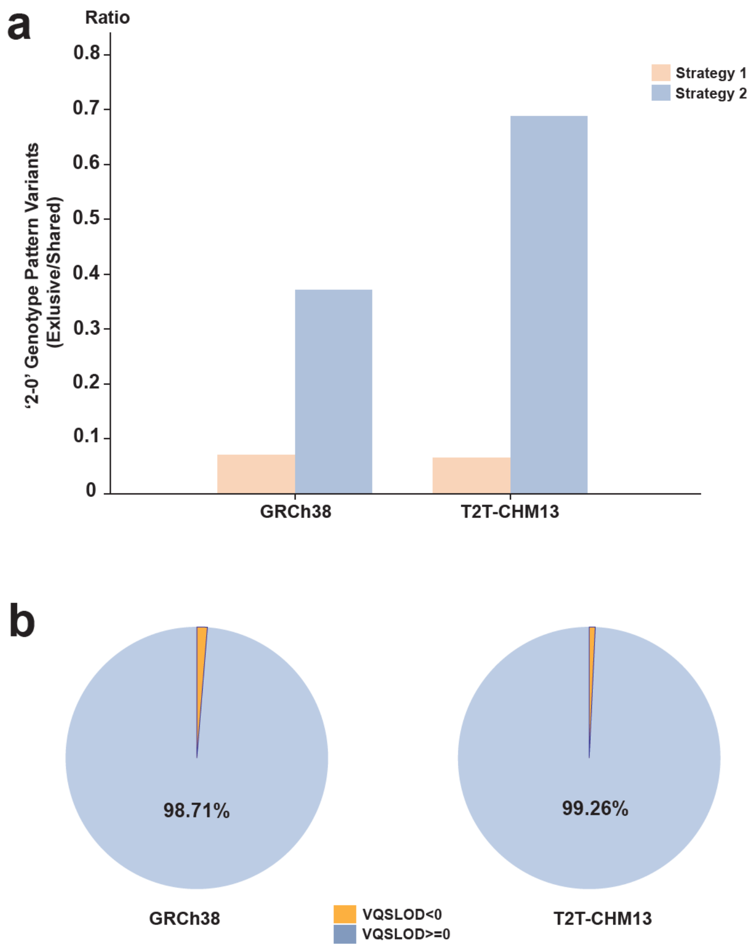


**Fig. S3. '2-0' genotype pattern variants under two pre-phasing strategies.** (a) The ratio of exclusive '2-0' genotype pattern variant count under each strategy and two-strategies-shared '2-0' genotype pattern variant count in GRCh38 and T2T-CHM13. (b) Pie charts show the ratio of the '2-0' genotype pattern variants caused by VQSLOD criterion of Strategy 2 in GRCh38 (left) and T2T-CHM13 (right).

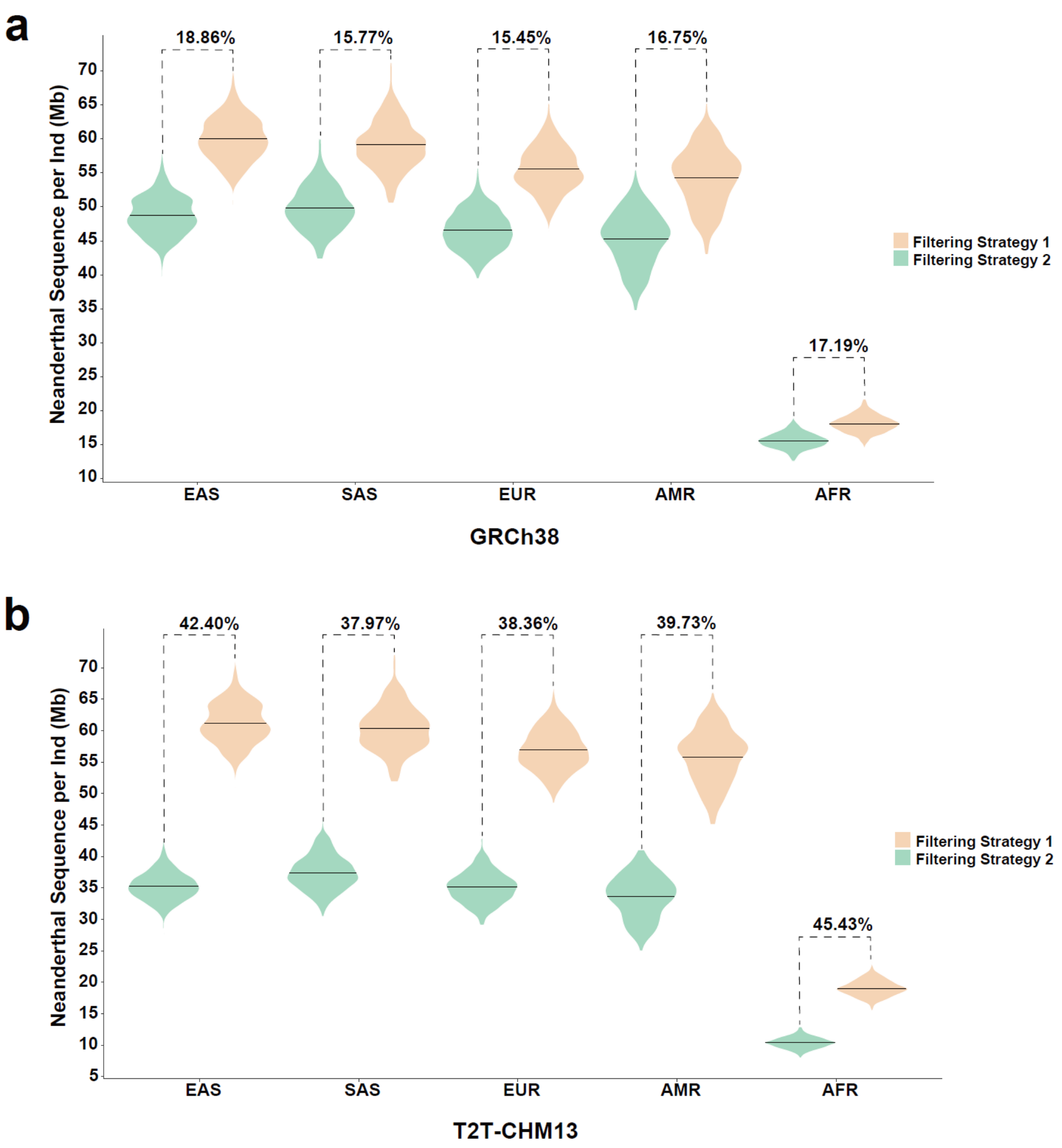


**Fig. S4. Comparison of Neanderthal introgression detected under different pre-phasing filtering strategies.** Violin plots show Neanderthal introgression bias per individual under two different pre-phasing filtering strategies in (a) GRCh38 and (b) T2T-CHM13.

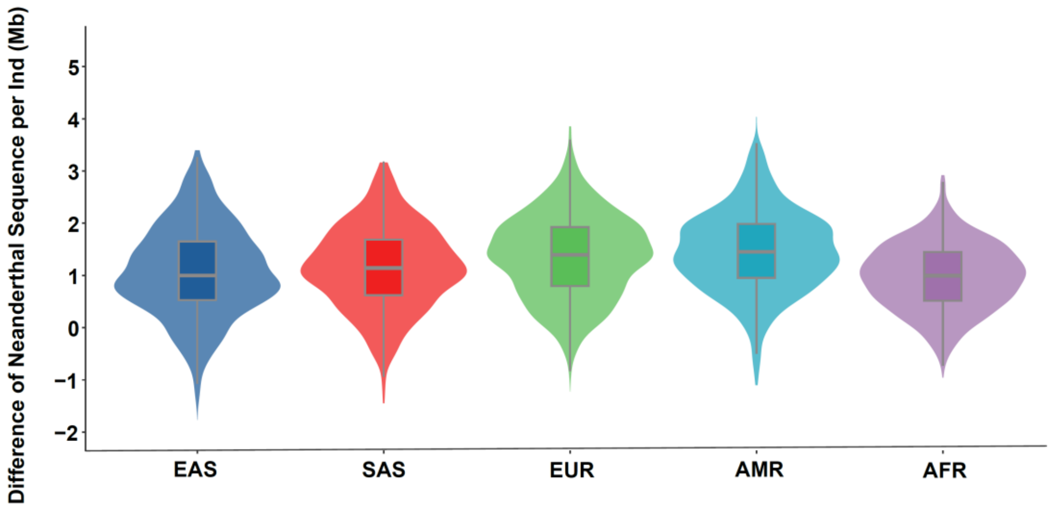


**Fig. S5. Difference of Neanderthal introgression per individual between T2T-CHM13 and GRCh38.** Violin plots show difference (Mb) of Neanderthal introgressed sequences per individual identified between T2T-CHM13 and GRCh38 in five populations from the 1000 Genomes Project (1KGP).
